# Dissecting cancer resistance to therapies with cell-type-specific dynamic logic models

**DOI:** 10.1101/094755

**Authors:** Federica Eduati, Victoria Doldàn-Martelli, Bertram Klinger, Thomas Cokelaer, Anja Sieber, Fiona Kogera, Mathurin Dorel, Mathew J Garnett, Nils Blüthgen, Julio Saez-Rodriguez

## Abstract

Therapies targeting specific molecular processes, in particular kinases, are major strategies to treat cancer. Genomic features are commonly used as biomarkers for drug sensitivity, but our ability to stratify patients based on these features is still limited. As response to kinase inhibitors is a dynamic process affecting largely signal transduction, we investigated the association between cell-specific dynamic signaling pathways and drug sensitivity. We measured 14 phosphoproteins under 43 different perturbed conditions (combination of 5 stimuli and 7 inhibitors) for 14 colorectal cancer cell-lines, and built cell-line-specific dynamic logic models of the underlying signaling network. Model parameters, representing pathway dynamics, were used as features to predict sensitivity to a panel of 27 drugs. This analysis revealed associations between cell-specific signaling pathways and drug sensitivity for 14 of the drugs, 9 of which have no genomic biomarker. Following one of these associations, we validated a drug combination predicted to overcome resistance to MEK inhibitors by co-blockade of GSK3. These results underscore the value of perturbation-based studies to find biomarkers and combination therapies complementing those based on a static genomic characterization.

## Introduction

Patient response to anticancer therapies is extremely variable and understanding the reasons for this variability is a major challenge in cancer research. One approach to address this problem is to identify biomarkers which correlate with therapy response. However, except for a few examples, no efficient biomarkers are available *(1)*. The problem of finding biomarkers has become even more important with the advent of targeted cancer therapies that are designed to affect specific molecular changes in cancer cells which drive the cancer, and which provide a larger number of treatment options. However, biomarkers that can be used to stratify patients for targeted drugs also remain largely elusive *(2).*

The most common approaches for patient stratification are currently based on genomic biomarkers, typically consisting of either expression or mutation of specific genes. The popularity of those biomarkers has been favoured by the advancements of sequencing technology and their subsequent decrease in cost, and, for some drugs, they have shown strong potential *(3–5)*. However, for many other cases no efficient genomic markers for patient stratification exist, and for those where markers exist, they have rather low power due to the complex nature of cancer. Furthermore, their actual clinical significance for precision oncology is questionable *(1)*.

Many modern drugs target signaling molecules, and elicit a response of the signaling network. Therefore, investigating the functioning of signaling pathways can provide new insights into these mechanisms and may help to unveil new therapeutic strategies and biomarkers *(6, 7)*. Understanding how drugs affect the signaling network as a whole is particularly important as signaling dynamic differs not only between tumors of different tissue, but also between tumors of the same tissue. For instance, for colon and liver cancer cell-lines, the dynamics of signaling networks have been shown to differ strongly between different cells of the same tissue of origin *(8–10)*.

Furthermore, models parameterized and experimentally calibrated using a specific cell-line have been used to define pathway dynamic biomarkers of therapeutic outcome, such as drug sensitivity *(11)* or patient survival *(12)*, based on model simulations in breast cancer and neuroblastoma, respectively.

The understanding of mechanisms of cellular response (network structure and dynamic behaviour), can be also useful to tackle the development of compensatory signaling mechanisms of drug resistance *(13)*, which is a recurrent problem for targeted therapy and is challenging to predict based only on genomic information. Signal transduction results from the integration of complex dynamic networks which can be modulated by mutations in oncogenes or tumor suppressors. The complex wiring confers a robustness to the cells that helps them to escape most single-agent- targeted treatments. Based on this idea, modeling of signaling pathways has been recently used to suggest combinatorial targeted therapies to effectively block multiple molecular pathways *(8, 14, 15)*: by integration of prior pathway knowledge with experimental observations cell-line-specific models were built, which could be used to simulate and thus prioritize possible combinatorial perturbation experiments.

In this paper, we investigate to what extent dynamic interactions between different signaling pathways play a role in characterizing the specific cellular responses to drugs and suggesting targeted combinatorial therapies. Furthermore we were interested to assess how these dynamic features fare against static genomic traits as markers of drug response. We do so by characterizing cell-type-specific models for a panel of 14 different colorectal cancer (CRC) cell-lines that are integrated with a large-scale drug screening *(16)*. We found associations with model parameters for 14 of the 27 drugs targeting our pathways of interests, for 9 of which there were no genomic biomarkers. These associations was used to define pathway dynamic biomarkers and interesting combinatorial therapies.

## Results

To identify biomarkers of drug response from dynamic logic models, we proceed as follows (**Fig. 1**). First, cell-line-specific models were built for 14 colorectal cancer (CRC) cell-lines. Each model is a set of logic ordinary differential equations *(17)* based on a prior knowledge network (PKN) manually curated from literature (**Fig.1A**) that is refined using CellNOpt *(18)* based on experimental data. Second, sensitivity data for our 14 cell-lines in response to drugs targeting nodes in the PKN or first neighbours were retrieved from the Genomics of Drug Sensitivity in Cancer (GDSC) panel *(16)* (**Fig. 1B**). Third, we investigated correlations between model parameters and drug response using Elastic Net *(19)* to select strong associations (**Fig. 1C**). All steps will be detailed in the following sections.

**Fig. 1.**
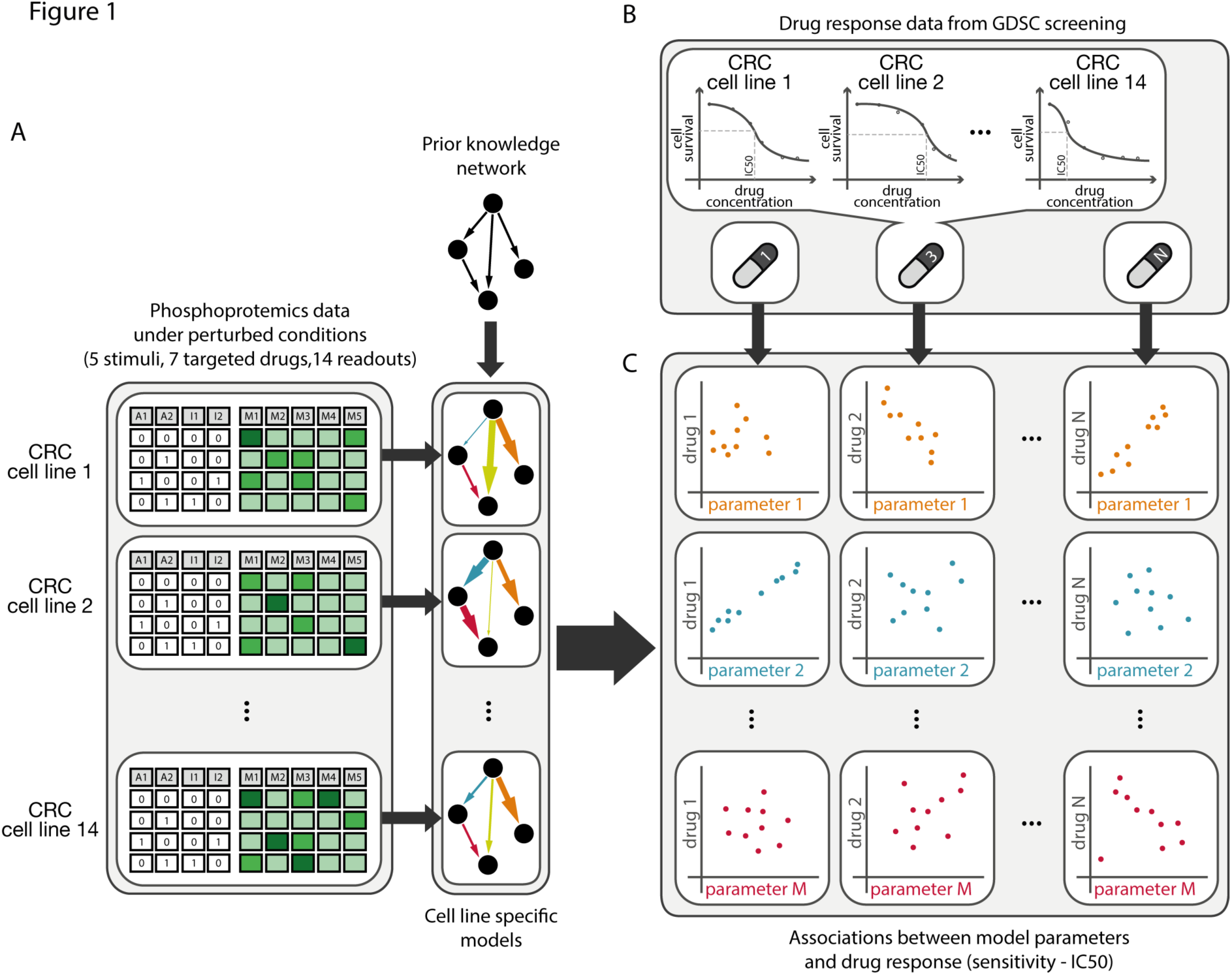
Schematic of the approach used to define biomarkers of drug response from pathway models. A. Specific predictive models for 14 colorectal cancer (CRC) cell-lines were built from perturbation data and prior information about network structure (PKN). B. Drug sensitivity data for the same 14 CRC cell-lines were retrieved from the Genomics of Drug Sensitivity in Cancer (GDSC) panel in response to 27 different drugs targeting the PKN or first neighbour. C. Elastic Net was used to select strong associations between model parameters and drug sensitivity.

### Strategy for model optimization: a toy example

Our starting point is a prior knowledge network (PKN), which is generic in the sense that it contains information about different cell types and is hence not clear about how many of the contained interactions are functional in a given cell type. To identify cell-specific functional components, the PKN was trained as a logic model to cell-specific data. The dynamics of the system were modelled using a formalism based on logic ordinary differential equations (ODEs) *(17)*, where a set of ODEs (one for each species in the model) was derived from the logic structure using a continuous update function (see **Materials and Methods**). Each species (i.e. node in the network) was characterized by a parameter *τ_i_* (with i=1,…,N where N is the number of species in the model) representing its life-time (*τ*=0 meaning that the node is not functional), and each regulatory interaction was defined by a sigmoid function, where a parameter k_i,j_ characterizes the strength of the regulation of species i dependent on species j (*k_i,j_*=0 meaning no regulation).

In order to be able to deal with large-scale networks including the effect of multiple signaling pathways and their cross-talks, building on *(18, 20)* we developed a new calibration approach that uses L1 regularization to prune the network by inducing sparsity thus reducing the size of the model. The approach can be illustrated using the *in silico* model in **Fig. 2A**, where the PKN includes 9 nodes and 14 edges. A subnetwork (10 edges shown in black in **Fig. 2A**) was used to generate *in silico* data for 2 readouts (nodes shown in blue in **Fig. 2A**), consisting of 9 perturbations, which are some of the possible combinations of 2 stimuli (green nodes) and 2 inhibitors (targeting red nodes). *In silico* data were generated for 10 random sets of parameters (one example shown in **Fig. 2B**) and were then used to optimize the model by minimizing *Q_LS_*(*τ,k*), which is defined as the sum of the squared difference between model predictions and true (*in silico*) values. Although the fit of the optimized models to the data was perfect, parameters could not be well estimated due to model redundancy and low identifiability. In order to improve parameter estimates, we redefined the objective function *Q* introducing a L1 regularization term on the parameters *τ_i_* to help the model remove the unconnected nodes (e.g. P4 and P5 in **Fig. 2A**). This was achieved by penalising the complexity of the model as:

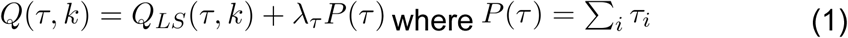

**Fig. 2.**
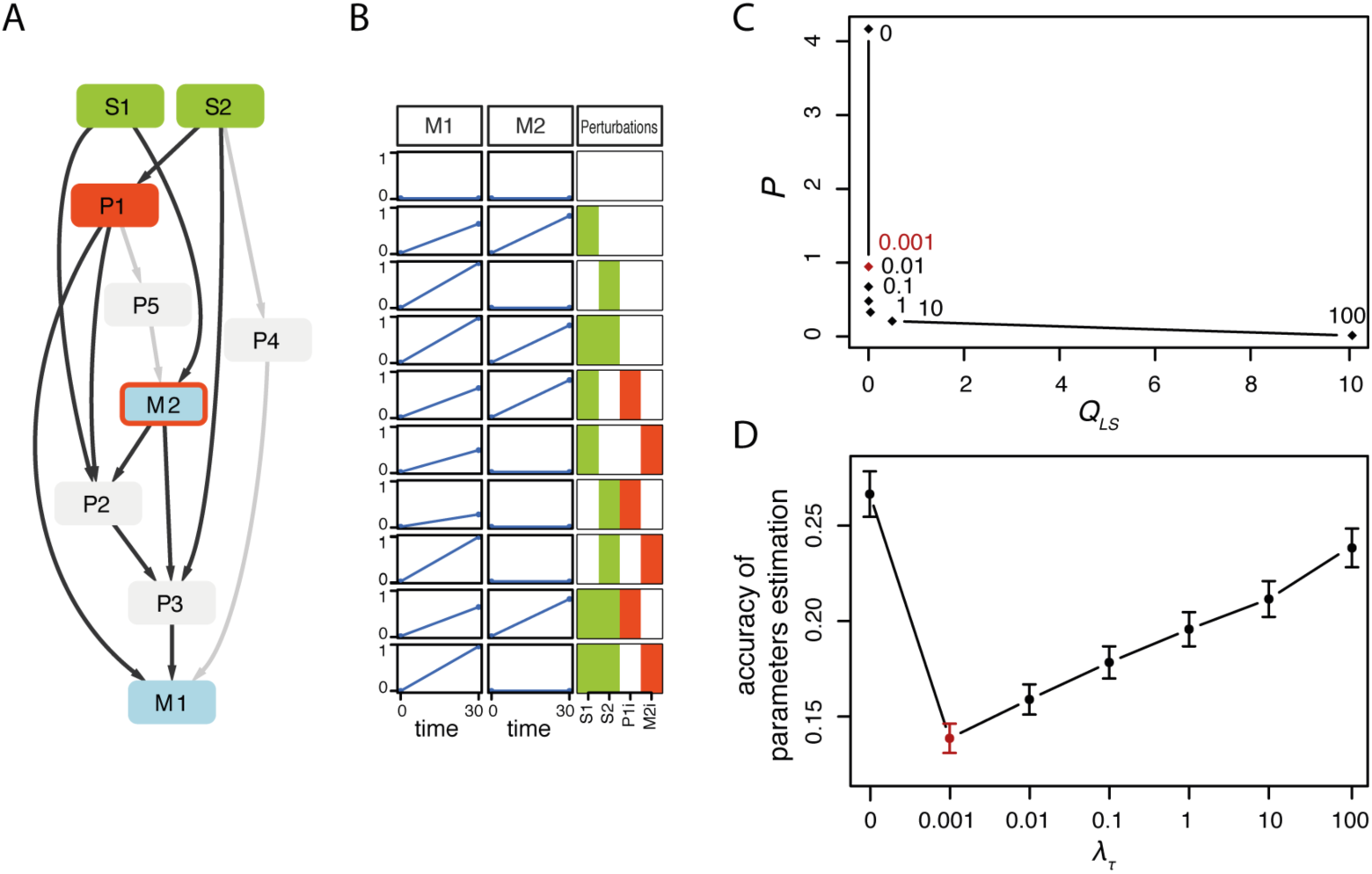
Toy example to illustrate modeling approach. A. Toy general network (PKN) with edges used for data simulation highlighted in dark grey. B. One (out of 10) example of simulated perturbation data used to illustrate L1 regularization. C. Goodness of fit (*Q_LS_*, i.e. sum of squared residuals) versus model sparsity (*P*, i.e. sum of estimated parameters) for different values of *λ_T_*, (λ_*T*_=0, 0.001, 0.01, 0.1, 10, 100) resulting in an L-shaped curve (each dot is the mean value obtained from the 10 *in silico* datasets). D. Mean accuracy (error bars represent standard deviation) of the estimated parameters (sum of the square of the difference between estimated and true parameters) as function of *λ_T_*. Optimal *λ_T_* is depicted in red.

The term λ_*τ*_ controls the importance of the regularization term. We plotted the effect of increasing values of λ_*τ*_ for the *in silico* model on *P* and *Q_LS_* (**Fig. 2C**), and on the accuracy of parameter estimates (as a sum of the square of the difference between estimated and true parameters) (**Fig. 2D**). As expected, for increasing values of λ_*τ*_, Q_LS_ tends to increase (worse fit to the data) while *P* tends to decrease (sparser model). Good values of λ_*τ*_ are those on the elbow of the L-shaped curve in **Fig. 2C**, corresponding to the best compromise between good fit and sparse model. Interestingly, these values (especially λ_*τ*_=0.001 which represents the most conservative choice in terms of regularization) also correspond to the best accuracy in estimating the parameters in **Fig. 2D**. Looking at the estimated parameters we could verify that for λ_*τ*_=0.001 the parameters *τ_P4_* and *τ_P5_* were indeed set to zero thanks to the regularization term in the objective function.

### Generation of cell-line-specific models

The previously illustrated approach was applied to systematically characterize how signaling pathways mediate differential cellular responses to drugs in colorectal cancer. For this purpose we performed a large-scale perturbation screening consisting of 14 phosphoproteins measured in 14 genetically heterogeneous colorectal cancer (CRC) cell-lines in 43 different perturbed conditions. This sums up to a total of 8428 different experimental data points which are represented in **Fig. 3** (experimental details in **Materials and Methods**). Perturbations consist of combinations of 7 kinase specific inhibitors (targeting IKK, MEK, PI3K, BRAF, TAK1, TBK1, mTOR) and 5 stimuli (EGF, HGF, IGF1, TGFb, TNFa).

**Fig. 3.**
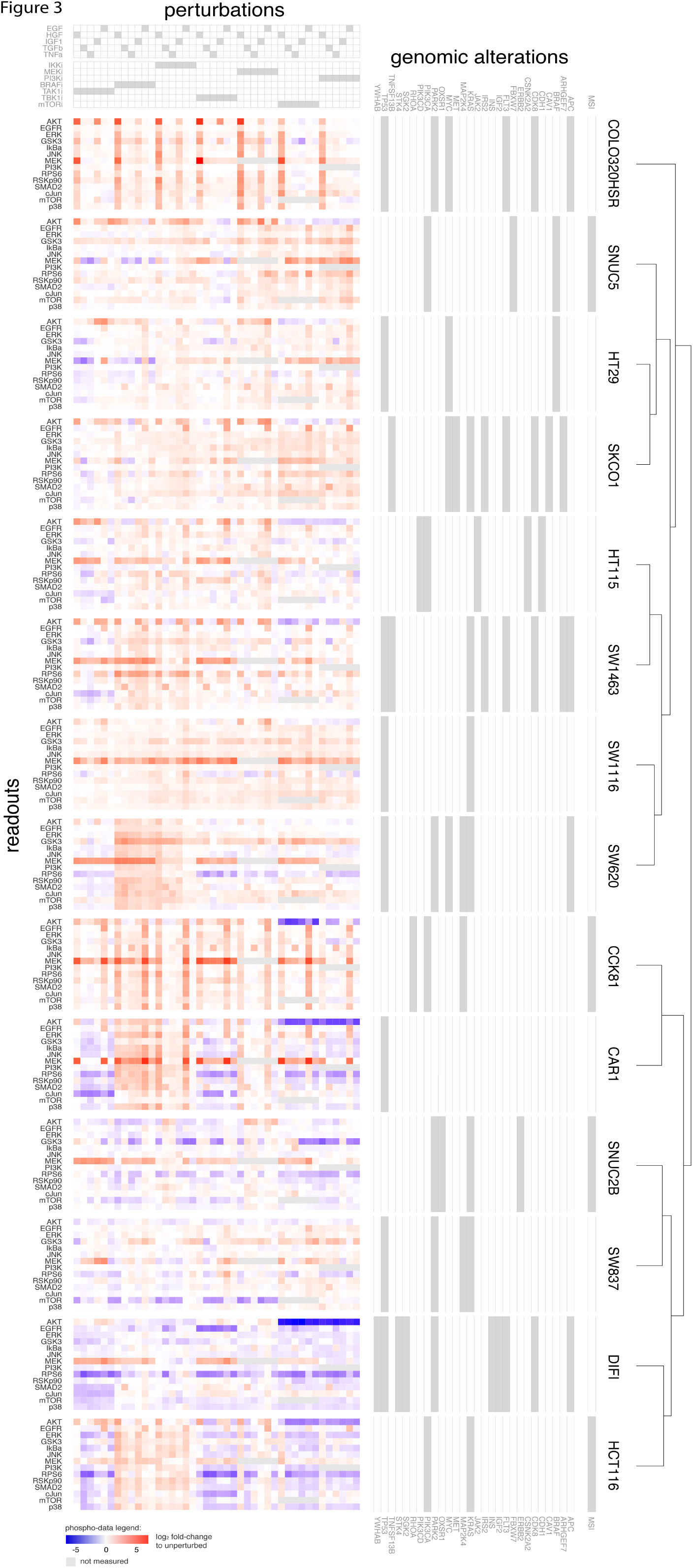
Large-scale experimental perturbation dataset. Data for 14 colorectal cancer cell-lines in response to 43 different perturbed conditions with 14 measured phosphoproteins. Genomic alterations affecting investigated pathways are also shown and cell-lines are clustered based on phosphorylation profile across all perturbations.

To characterize the signaling pathways we started from a comprehensive general prior-knowledge network (PKN) derived from the literature and public databases using OmniPath *(21)*. The PKN was first compressed to reduce model complexity without affecting logic consistency as described in *(22)* (complete PKN in **fig. S1**), while maintaining interactions between the measured nodes (in blue in **Fig. 4A**), the inhibited nodes (in red) and the simulated nodes (in green). The definition of the general model also took into consideration that one of the kinase inhibitors, PLX4720, works as selective BRAF inhibitor in cell-lines where BRAF is mutated in V600E (i.e. HT29 and SNUC5 in our panel), but induces a paradoxical activation of wild type BRAF cells, therefore-we modeled it as a stimulus for those cell-lines.

**Fig. 4.**
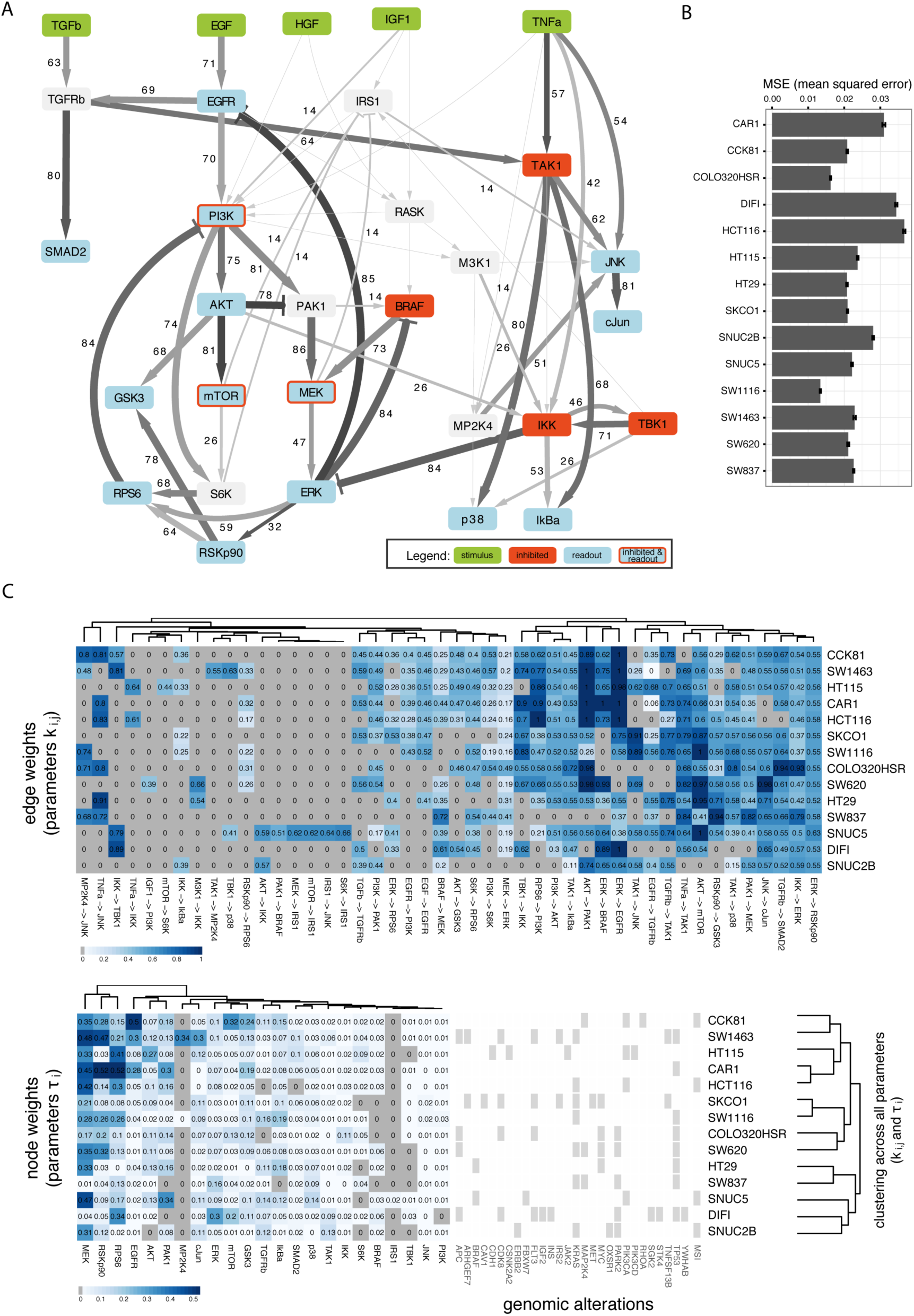
Results of cell type specific model optimization. A. Compressed PKN with edge width and values representing the level of heterogeneity of the corresponding model parameter among cell-lines (percentage of pairwise tests between cell-lines for which null hypothesis of equal distribution was rejected) and edge colour representing median value of the corresponding parameter across all cell-lines. B. Mean squared error (MSE) for each cell-line, error bars represent standard deviation derived from bootstrap. C. Estimated parameters *k_ij_* (edge weight) and *τ_i_* (node weight) which are different from zero in at least one cell-line. Clustering on the right is based on all estimated model parameters which does not correlate well with genomic alterations.

The following three-step optimization procedure was used for each cell-line (see **Materials and Methods** for more details). First, L1 regularization was applied to parameters *τ_i_* to remove unconnected nodes, as described in Equation 1. Secondly, values of *τ_i_* were fixed to the values estimated in Step 1 (shown in **Fig. 4C**) and optimization was repeated applying this time the L1 regularization to parameters *k_ij_* to induce sparsity in the network. Thirdly, bootstrap distribution was then obtained for parameters *k_ij_* by repeating the optimization on resampled experimental data with replacement (150 times). Median values for the *k_ij_* parameters estimated at Step 3 are shown in **Fig. 4C** for all cell-lines. The resulting models were comparably sparse across cell-lines, with a percentage of values set to zero ranging between 12–28% for parameters *τ_i_* and 51–73% for parameters *k_ij_*. The optimized models showed good fit to the experimental data for all cell-lines, with mean squared error (MSE) within the range [0.013, 0.036] as shown in **Fig. 4B**.

### Heterogeneity of the cell-line specific models

We further analysed the heterogeneity of the estimated parameters *k_ij_*, in order to understand the variability in signal processing across the cell-lines. For each parameter, distributions across all cell-lines were compared using Kruskal-Wallis rank sum test (one-way analysis of variance on ranks) to test if estimated parameters (from bootstrap) stem from the same distribution for all cell-lines. For 46 out of 63 parameters the null hypothesis was rejected (p-value<0.05 after Bonferroni correction, and large effect size *w* > 0.5, see **Materials and Methods**), meaning that bootstrapped parameters come from a different distribution for at least one cell-line. The 17 Parameters for which the null hypothesis of equal mean rank was accepted showed a median value equal to zero for all cell-lines. For the remaining 46 parameters, post hoc pairwise tests (91 possible combination of 14 cell-lines) were performed using Wilcoxon rank sum test to additionally explore how many cell-lines have different parameter distributions (i.e. p-values Bonferroni corrected for multiple hypothesis testing < 0.05 and high *r* effect size > 0.5, see Online Methods). The resulting percentages are shown in **Fig. 4A** as numbers next to each corresponding edge (only for the 46 edges passing the one-way analysis of variance test) and mapped as width of the edge, while edge colour intensity maps its mean value across cell-lines. As we might expect, genetic alterations alone are not representative of clusters of cell-lines neither at the level of phosphorylation response (**Fig. 3**) nor at the level of pathway models (**Fig. 4C**).

Among the 46 parameters *k_ij_* showing different distributions across cell-lines, only 34 were different from zero in at least three cell-lines. For further analysis, we defined these parameters as highly heterogeneous. As for parameters *T_i_* only 1 out of 25 parameters was set to zero across all cell-lines and 21 parameters were different from zero in at least three cell-lines. Among these, 12 were defined as highly heterogeneous, showing reasonable variability across cell-lines (standard deviation > 0.04). Although we are aware that the negative interaction between IKK and ERK might be artificial due to unspecific effect of the used IKK inhibitor, which causes an increase in ERK phosphorylation *(8)*, we decided to keep the interaction in the model as we are interested in capturing all differential responses between cell-lines. From our analysis, we conclude that cell-lines are very different and hence it is important to characterize each cell-line with a specific signaling model.

### Association of model parameters with drug response data

We next investigated if the previously defined highly heterogenous model parameters can be associated with the efficacy of drugs. We reasoned that since the mode of action of multiple drugs involves affecting the functioning of the pathways we were modeling, their functional status should affect the efficacy of the drugs.Since all our 14 colorectal cancer cell-lines are part of the Genomics of Drug Sensitivity in Cancer (GDSC) panel *(16)*, we could make use of the drug sensitivity data included in this comprehensive dataset in response to a large panel of drugs. In particular, in order to focus on biologically relevant associations, we selected only those drugs targeting nodes in our network (PKN) or first neighbours, i.e. targets that directly regulate a node in the PKN based on Omnipath *(21)*, which is a curated ensemble of multiple pathway resources. We additionally excluded drugs showing a low variability in the response across our 14 cell-lines (less than three sensitive and three resistant cell-lines based on the classification used in *(16)*) and those for which sensitivity data were missing for more than three of our 14 cell-lines. For the remaining 27 drugs, we investigated the association with our model parameters using cross-validated Elastic Net (details in **Materials and Methods**), using IC50s (half maximal inhibitory concentration) as a measure of drug sensitivity. We found a total of 146 robust associations (out of the 1242 possible ones), with 14 of our 27 drugs being explained by at least one model parameter. Top 50 associations are shown in **Fig. 5A** and all associations are shown in **fig. S2**. Resulting associations were then compared with those between drugs and genomic alterations (functional mutations and copy number alterations) acting on PKN nodes or first neighbor. Genetic biomarkers were identified using ANOVA as described in *(16)* (without correction for multiple hypothesis testing due to too large number of comparisons, see **fig. S3**) resulting in associations for 12 of our 27 drugs, five of which also showed associations with model parameters (corresponding genetic biomarkers are reported for the drugs in **Fig. 5A**).

**Fig. 5.**
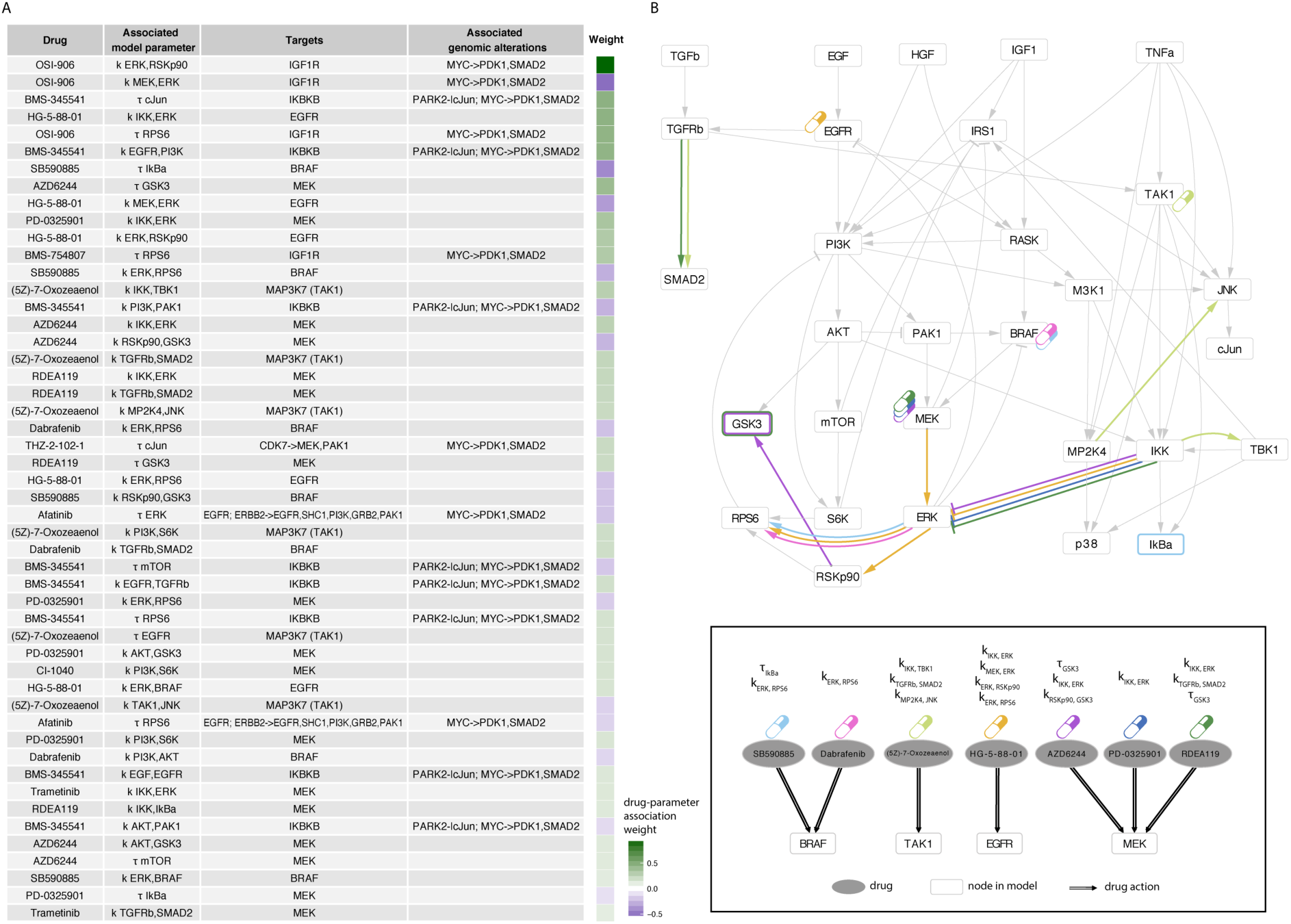
Associations between model parameters and drug sensitivity. A. Top 50 associations resulting from Elastic Net selection between a drug (first column) and a model parameter (second column). For each drug, the corresponding targets are shown in third column and the associated genomic biomarker (if any) in the fourth column. Analysis is limited to drugs and mutations acting on PKN or first neighbour. For both drugs and mutations, corresponding PKN targets are indicated with an arrow (-> for positive regulation; -| for negative regulation) if mutations and/or drug targets are first neighbour regulating nodes in the PKN (which are named after the arrow). B. Among the top 25 parameter-drug associations, those involving drugs with no genetic biomarkers are schematically mapped to the signaling pathways. Drug targets for the 7 drugs listed as ellipses in the bottom panel, are shown in the compressed PKN (upper panel) using the corresponding coloured drug symbols. Corresponding model parameter associations for each drug are shown in the PKN using the same colour code to mark edges corresponding to regulator parameters k_ij_ (edge from *i* to *j*) and nodes border for corresponding life-time parameters *T_i_* (node *i*).

### Interpretation and validation of the dynamic biomarkers

We then focused on the 9 drugs whose efficacy was not significantly associated with mutations or copy number alterations but instead with pathway biomarkers, to explore how our approach provides insights in otherwise unexplained drug efficacy. Top associations are illustrated in **Fig. 5B**, visualized on the network (PKN) to show, for each drug, both the target(s) and the associated pathway parameter(s).

For example, multiple parameters related with MEK-ERK pathway showed association with EGFR signaling inhibitor (HG-5-88-01) and with BRAF inhibitors (SB590885 and Dabrafenib). Interestingly, many recent studies reported synergistic effects when combining MEK inhibitors with EGFR inhibitors *(8, 23, 24)* or with BRAF inhibitors *(8, 25, 26)* in different cancer types. In particular, focusing on colorectal cancer, combination of EGFR and MEK inhibitors was suggested to overcome resistance to MEK inhibitors in BRAF and KRAS mutants *(8, 27)* and it was suggested in *(24)* as potential therapy for CRC patients who become refractory to anti-EGFR therapies. A significant increase in response was also shown on a phase I study for CRC patients treated with a combination of three drugs targeting MEK, BRAF and EGFR respectively, in BRAF V600E mutants *(28)* and is currently in Phase I/II study (NCT01750918 in clinicaltrials.gov). These examples underline the potential of our approach in suggesting targeted combinatorial therapies.

Another interesting example is the association of different MEK inhibitors (AZD6244, RDEA119, Trametinib, PD-0325901, CI-1040) with multiple model parameters related to GSK3 (*τ_GSK3_*, *K_RSKp90,GSK3_*, *K_AKT,GSK3_*), as shown in **Fig. 6A-C**, with each parameter being associated with at least two of the inhibitors. The top association among these (which is overall ranked 8^th^ in **Fig. 5A**), between AZD6244 and *τ_GSK3_*, is shown in the centre of **Fig. 6D**. Our model predictions suggest that the combination with a GSK3 inhibitor could improve the sensitivity for cell-lines resistant to MEK inhibitors (i.e. high IC50) and with functional GSK3 (i.e. high *T_GSK3_*. On the contrary, cell-lines with nonfunctional GSK3 (i.e. low *τ_GSK3_*) should not be affected by GSK3 inhibitors. In order to experimentally test this hypothesis, we combined a MEK inhibitor (Trametinib) with two GSK3 inhibitors (SB216763 and CHIR-99021) at increasing concentration (see **Materials and Methods**) as shown in the external plots in **Fig. 6D**. As expected, cell-lines with nonfunctional GSK3, based on the corresponding model parameter, do not show an increased sensitivity when treated in combination with any of the two tested GSK3 inhibitors, regardless of their response to MEK inhibitors (e.g. HT29 and SNUC2B for a sensitive and a more resistant case). On the contrary, cell-lines with more functional GSK3 tend to show an improved sensitivity under the combinatorial treatment (see HT115, SW1116, SW1463 and CCK81) even at the low drug concentrations tested.

**Fig. 6.**
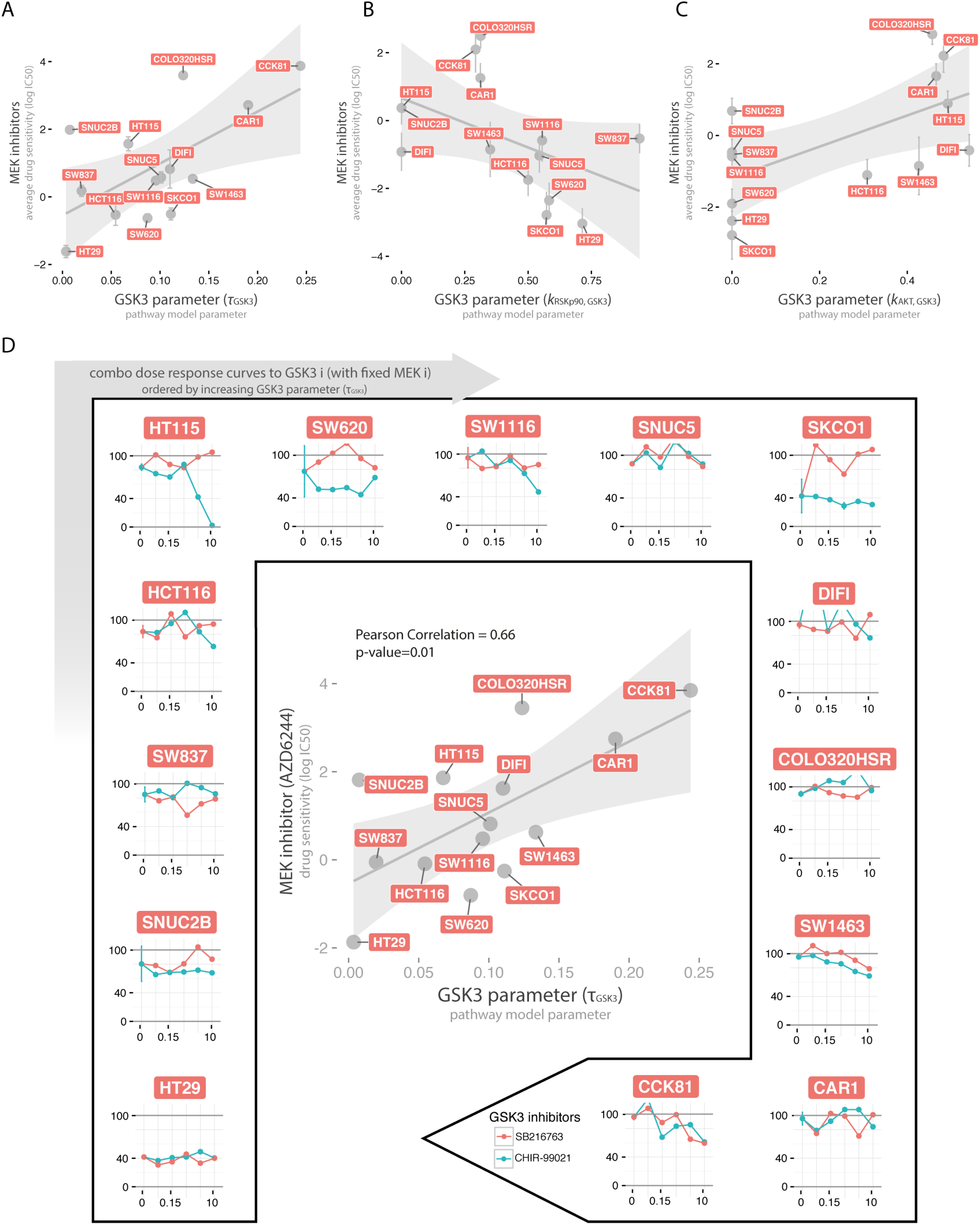
Association between MEK inhibitors and GSK3 model parameters. A-C. Scatterplots of the three GSK3 related model parameter (*τ_GSK3_*, *k_RSKp90_*, *k_AKT,GSK3_* in panel A, B and C respectively) versus the mean sensitivity of the respective associated MEK inhibitors (at least two for each parameter), with error bars representing the standard error. D. In the centre, scatterplot of the strongest association between MEK inhibitor (AZD6244) and GSK model parameter (*τ_GSK3_*). In the external plots, dose-response curves when combining MEK inhibitor (Trametinib) (at fixed concentration 0.005 μM) with two GSK3 inhibitors (SB216763 and CHIR-99021) at increasing concentrations (0, 0.039, 0.156, 0.625, 2.5, 10 μM). Plots for the 14 CRC cell-lines are ordered clockwise according to their *τ_GSK3_* parameter value.

## Discussion

While it is broadly accepted that alterations in the functionality of signaling pathways largely determine the efficacy of kinase inhibitors used in the clinic, a complete understanding of their relationship is lacking. In this study we investigate this relationship and we show that the dynamic of signaling pathways can determine the efficacy of targeted drug treatments, in particular in cases where genomic data cannot. Additionally, we show that this information can be used to guide combinatorial therapies.

Our means towards this end were cell-specific mechanistic models of signal transduction for 14 colorectal cancer cell-lines trained with dedicated phosphoproteomic data upon perturbations. We built our models based on differential equations to capture the continuous aspects of signal strength and time-dynamics, but using a logic formalism that allowed for straightforward interpretability of the model parameters in terms of life-time of each species in the network and strength of the regulatory interactions. We could then study how these model parameters, which determine the dynamic behaviour of the pathway, are related with the global cellular sensitivity to cancer drugs. We found some strong correlations between model parameters and drug sensitivity, such as between GSK3 life-time and MEK inhibitors, that suggest that the functionality of pathway interactions is indeed related to the efficacy of the drugs.

A key aspect of our study is the opportunity to analyze the heterogeneity of dynamic functionality (which is described by the model parameters) of signal transduction among a large panel of colorectal cancer cell-lines and how it relates to therapeutic drug response. The investigation of the heterogeneity of signaling pathways was empowered by the uniqueness of our dataset, consisting in 14 phospho-proteins measured under 43 differently perturbed conditions, which allowed us to characterize cell-line-specific models for 14 cell-lines derived from the same tissue of origin. We found that about half of the model parameters are highly heterogeneous in our panel of cell-lines and that heterogeneity cannot be explained by genomic alterations alone. Since these findings were obtained with a relatively limited sampling of the signaling status, we expect that many more associations can be found with a broader characterization. Mass-spectrometry based approaches, while still more costly and laborious, are under active development and likely to enable such an analysis in the near future.

Associations between model parameters and drug sensitivity were used to define interesting points of intervention to overcome resistance to specific drugs in specific cell-lines, thus paving the way for personalized combinatorial therapies. Some of the combinatorial therapies suggested using this approach, like the combination of MEK inhibitors with EGFR and/or BRAF inhibitors, are supported by literature (as reported in the Results) and are currently in clinical trials. We could also suggest novel combinatorial interventions, like the combination of MEK and GSK3 inhibitors, that we validated experimentally. Interestingly, the inhibition of GSK3 showed promising anticancer effect in recent studies on cancer cell-lines *(29, 30)* and its potential interaction with MEK-ERK pathway has also been suggested in this context *(30, 31)* but, as far as we know, no proven synergistic effect had previously been reported.

Our findings underscore the value of studying signaling pathways dynamic to better understand tumor phenotypes and to exploit this knowledge to suggest new therapeutic strategies *(32)*. Such an approach is fundamentally different from the currently common characterization of various ‘omics’ layers at the basal level and it can be exploited to prevent the dynamic adaptation mechanisms underlying drug resistance *(14)*. Accordingly, we were able to find various pathway dynamic biomarkers for drugs with no genetic biomarkers. Hence, we believe that a perturbation-based strategy, even if restricted to fewer genetic backgrounds and only monitoring selected proteins, can provide a complementary strategy for biomarker discovery in cancer and beyond.

## Materials and methods

### Definition of logic ordinary differential equations

The logic ordinary differential equation formalism *(17)* is based on ordinary differential equations derived from logic models using a continuous update function *B_i_* for each species *x_i_*, which can assume continuous values {0, 1}. The differential equation for species *i* is defined as follow:

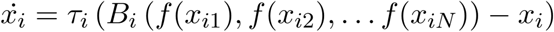

Where *τ_i_* is the life-time of species *i*.*τ_i_* = 0 means that the node is disconnected by the rest of the network and higher values of *τ* represent faster response, that can be interpreted as more functional species, *x*_*i*1_,*x*_*i*2_…, *x*_*iN*_ are the N regulators of *x_i_* and each regulation is defined by a transfer function *f*(*x_ij_*) with parameters *𝑛_ij_* and *k_ij_*:

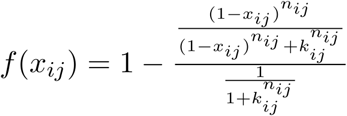

Where parameters *n_ij_* were fixed to 3 and parameters *k_ij_* were optimized (constrained between 0 and 1 included). As shown in **fig. S4**, *k_ij_* = 0 means that the transfer function is equal to 0 for any value of the regulator j and can be interpreted as the edge j→i is not functional. Higher values of *k_ij_* represent increasing strength of the regulatory interaction. For consistency with Boolean models (see *(17)* for more details), the transfer function is defined to be *f*(*x_ij_*) = 1 for *x_ij_* = 1 (except for the *k_ij_* = 0 case)

Optimization of parameters *τ_i_* and *k_ij_* was performed using MEIGO *(33)* and CNRode add-on of the CellNOptR package *(18)* using a modified version of the objective function to include the parameter regularization described in the paper, as well as an additional penalty to promote that the simulated values at t=30 min (corresponding to the measured time point) are at steady state in order to match the experimental assumptions.

### Experimental setup of perturbation screen

Human colorectal cell-lines (CAR1, CCK81, COLO-320-HSR, DIFI, HCT116, HT115, HT29, SK-CO-1, SNU-C5, SW1116, SW1463, SW620 and SW837) are described in the COSMIC cell-lines project database (cancer.sanger.ac.uk/cell_lines). All cell-lines were cultured in either RPMI or DMEM/F12 medium with 10% FBS and were incubated at 37°C and 5% CO_2_. Before perturbation commenced cells were starved overnight in serum free medium. At 90 minutes before lysis the cells were treated with inhibitors (or solvent control DMSO) and at 30 minutes before lysis cells were stimulated with ligand (or solvent control H2O). This procedure was conducted for all single and combined inhibitor-ligand perturbations. The following inhibitors were used: MEKi AZD6244 (4μM), PI3Ki AZD6482 (10μM), mTORi AZD8055 (2μM), TBK1 i BX-795 (10μM), IKKi BMS-345541 (25μM), BRAFi PLX4720 (5μM) and TAK1 i 5Z-7-Oxozeanol (5μM). Concentrations were selected to inhibit target while minimizing off-target activity. We further used the following ligands: EGF (25ng/ml), HGF (50ng/ml), IGF1 (10ng/ml), TGFb (5ng/ml) and TNFa (10ng/ml). After treatment and incubation, lysates were collected and analyzed with the BioPlex Protein Array system (BioRad, Hercules, CA) as described earlier *(8)* using magnetic beads specific for AKT^S473^, c-Jun^S63^, EGFR^Y1068^, ERK1/2^T202 Y204/T185,Y187^, GSK3A/B^S21/S9^, IkBa^S32,S36^, JNK^T183,Y185^, MEK1^S217,S221^, mTOR^S2448^, p38^T180,Y182^, PI3K^Y458^, RPS6^S235,S236^, p90RSK^S380^ and SMAD2^S465,S467^. The beads and detection antibodies were diluted 1:3. For data acquisition, the BioPlex Manager software and R package lxb was used.

### Drug combination experiments

Experiments were performed using MEKi Trametinib (0.005μM) as anchor drug and testing its effect on our 14 colorectal cancer cell-lines alone and in combination with 5 concentrations (from 10μM with 4-fold dilutions: 0.039μM, 0.156μM, 0.625μM, 2.5μM, 10μM) of two GSKi (SB 216763 and CHIR-99021). For available replicates median value was computed. Dose-response curves were normalised between 0 and 100 with 100 corresponding to the negative control (cells with no treatment) and 0 to the positive control (cells treated with Staurosporine 2μM). Same DMSO volume was maintained for all experiments (negative and positive controls, single drug and drug combination). Cell number was measured after 3 days of constant drug exposure using CellTiter-Glo reagent as described by the manufacturer (Promega).

### Data preprocessing

For each cell-line, data were processed separately for each of the 14 measured phosphoproteins. The value of the control (unperturbed condition) was estimated as the median value of the 4 replicates (to decrease the risk of bias) and log2 fold changes with respect to the control were then computed for each of the 42 perturbed conditions. As required by our logic formalism, data were then linearly scaled between 0 and 1, with 0.5 corresponding to the basal (control) condition.

### Model optimization

The optimization procedure applied to each cell-line is the following:

1. L1 regularization was applied to parameters *τ_i_* to remove unconnected nodes, as described in Equation 1, for five increasing values of λ_*T*_ (λ_*T*_=0, 0.01, 0.1, 1, 10). To increase the chances to obtain good solutions, 10 independent runs of the optimization were run in parallel. The best value for λ_*T*_ was selected (λ_*T*_=0.1) which provided the optimal balance between fitting precision and network size as described in the example, using the L-shaped curves (**fig.S5**).
2. Values of *τ_i_* were fixed to the values estimated in Step 1 for λ_*T*_=0.1 and optimization was repeated applying this time the L1 regularization to parameters *k* to induce sparsity in the network, with objective function defined as:

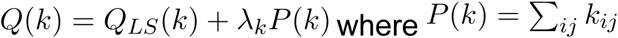 As in step 1, optimization was repeated for increasing values of λ_*k*_ (λ_*k*_=0, 0.01, 0.1, 1, 10) with 10 independent runs each, and best λ_*k*_ was selected based on L-shaped curves (**fig. S6**) to λ_*k*_=0.1.
3. Bootstrap distribution was obtained for parameters *k_ij_* (for λ_*k*_=0.1 from Step 2) repeating the optimization by resampling experimental data 150 times with replacement.

### Elastic net

In order to select only robust features, Elastic net training was iterated *M* times (where *M* is the total number of observations), each time leaving out one observation (sampling without replacement). For each iteration, leave-one-out cross-validation was applied to tune Elastic Net hyperparameters. Median values were computed for the weights of the features across the *M* iterations, in order to select only features appearing in at least half of the iterations.

### Statistical Analysis

ANOVA was used to define genomic markers of drug sensitivity using the GDSC tools (pypi.python.org/pypi/gdsctools/0.2.0). Microsatellite instability was used as covariate and threshold on minimum size of the positive and negative population for each feature set to 3. The threshold for significance of the p-value was set to 0.05, but no correction for multiple hypothesis testing was applied.

Kruskal-Wallis rank sum test (which is a one-way analysis of variance on ranks) was used to test if estimated parameters (from bootstrap) derive from the same distribution for all 14 cell-lines (null hypothesis rejected if different for at least one group). Effect size *w* was computed as 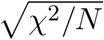 where X^2^ is the statistics from the test and *N* is the number of observations. Effect size >0.5 is considered as large effect. P-values were Bonferroni corrected and threshold was set to 0.05. Wilcoxon rank sum test was then used for post hoc pairwise test on parameters passing the Kruskal-Wallis rank sum test, using as effect size 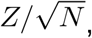 where *Z* is the statistics from the test and *N* is the number of observations. Effect size >0.5 is considered as large effect. P-values were Bonferroni corrected and threshold was set to 0.05. Rank type test were preferred over parametric tests because they are highly robust against non-normality.

## Supplementary Materials

Fig. S1. Extended version of the prior-knowledge network (PKN).

Fig. S2. Associations between model parameters and drug sensitivity.

Fig. S3. Genomic biomarkers of drug sensitivity.

Fig. S4. Transfer function used in logic ordinary differential equations.

Fig. S5. L-shaped curves for *τ* parameters.

Fig. S6. L-shaped curves for *k* parameters.

Data file S1 Optimised cell-line-specific models

Data file S2-S15 Phosphorylation perturbation data for the 14 cell-lines (MIDAS format)

## Acknowledgments

We thank members of the Saez laboratory for useful feedback, especially Luz Garcia-Alonso for help on mutations and Attila Gabor for fruitful discussion on model definition and David Henriques for support with CNORode. We thank Howard Lightfoot for help and assistance with drug combination data.

## Author contributions

F.E. designed and performed the computational analysis and wrote the manuscript. V.D.M. and T.C. contributed to the computational analysis. B.K. Contributed to experimental design, the interpretation of the results and participated in writing the manuscript. A.S. and F.K. generated the experimental data. M.D. contributed to data preparation and analysis. M.G. Contributed to experimental design and experimental supervision. N.B. Supervised experiments, contributed to interpretation of the results, and participated in writing the manuscript. J.S.R Supervised the computational analysis and the overall project, and participated in writing the manuscript. M.G., N.B., and J.S.R. conceived the project.

